# DS5: An Open-Source Framework for Standardized High-Throughput Drug Screening Data Storage, Analysis, and Drug Prioritization

**DOI:** 10.64898/2026.09.22.753533

**Authors:** Huiyi Yang, James Lubkowitz, Gabor Marth, Samuel Cheshier, Chieh-Hsiang Yang, Alana Welm, Philip J. Moos, Xiaomeng Huang, Yi Qiao

**Affiliations:** Department of Biomedical Informatics, University of Utah, Salt Lake City, UT, USA; Department of Human Genetics, University of Utah, Salt Lake City, UT, USA; Huntsman Cancer Institute, University of Utah Health, Salt Lake City, UT, USA; Primary Children’s Hospital, Intermountain Health & University of Utah School of Medicine, Salt Lake City, UT, USA; Department of Oncological Sciences, University of Utah, Salt Lake City, UT, USA; Department of Pharmacology and Toxicology, University of Utah, Salt Lake City, UT, USA

## Abstract

**Background:** High-throughput drug screening (HTS) is increasingly used in precision oncology across cell lines, patient-derived models such as organoids (PDO) and xenograft organoids (PDxO), and *ex vivo* patient samples, with multiple active clinical trials incorporating HTS for individualizing treatment. Yet HTS efforts predominantly rely on *ad hoc*, spreadsheet-based workflows, with no standardized infrastructure for storing, analyzing, and comparing screens across patients, cohorts, or institutions — limiting reproducibility and slowing clinical data exchange. Existing tools address isolated pieces of the workflow, handling either data management or analysis and visualization, but none integrate standardized storage, reproducible dose-response analysis, individualized drug prioritization, and clinician and molecular tumor board friendly reporting in a single open-source framework.

**Results:** We present **DS5**, an open-source Python framework that operates on an HDF5-based *DS5* file format that unifies standardized storage, quality control, dose-response analysis, cohort-level drug prioritization, and automated reporting in a single lightweight package. DS5 enforces raw-data immutability, integrates RxNorm-based drug name standardization, and scales from small clinical cohorts to large pharmacogenomics datasets on a standard laptop. We validated DS5 against two public benchmarks spanning over 428,000 drug–cell line pairs, recovering published LN_IC50 values from GDSC2 (Pearson *r* = 0.973) and Emax values from CTRP (*r* = 0.961). Using a published breast cancer PDxO dataset (45 compounds × 16 models), we reproduced GR-based profiles via a custom-metric extension and used DS5’s cohort-level prioritization to predict *in vivo* tumor response with ROC AUC = 0.907 across 30 drug–PDX pairs. Finally, we demonstrate the utilities of DS5 in an ongoing pediatric brain tumor precision oncology initiative, where its cohort-normalized drug rankings and automated reports directly support molecular tumor board discussions.

**Conclusion:** DS5 provides the cancer basic and translational research community with a standardized file format and reproducible computational analysis methods to support functional drug screening experiments; and its reporting features facilitate bridging raw drug screening measurements to translational decisions. DS5 is released open-source under the permissible MIT license.

## Introduction

High-throughput drug screening (HTS), the process of applying panels of drugs at multiple concentrations to biological samples and measuring response, is increasingly used in functional precision oncology across a range of model systems, including cell lines, patient-derived models such as organoids (PDO) and xenograft organoids (PDxO), and *ex vivo* patient samples (1–4).

The fundamental premise of functional precision oncology is that drug sensitivity measured directly in a patient’s own tumor cells can inform treatment selection by capturing the integrated functional consequences of a tumor’s complex molecular landscape, including epigenetic states, protein expression levels, and microenvironmental factors that are not fully reflected in molecular profiling data (5). Typically, patient or patient-derived cells are plated in a microwell format (96- or 384-well plates), exposed to compound libraries at multiple concentrations for 48–72 hours, and cell viability is quantified using luminescence-based readouts such as the CellTiter-Glo (6) assay. Multiple active clinical trials now incorporate HTS to guide individualized treatment decisions (2,7), and the approach has demonstrated clinical feasibility in both pediatric and adult cancers (1). The prospective SMARTrial demonstrated that *ex vivo* drug response profiling is clinically feasible in hematologic malignancies and that *ex vivo* chemoresistance predicts *in vivo* chemotherapy failure (7). In pediatric cancers, the Australian ZERO/TARGET pilot study integrated *in vitro* high-throughput drug screening and *in vivo* PDX testing alongside genomic profiling for 56 high-risk patients, identifying treatment options in 70% of cases and leading to therapy changes with clinical benefit in a subset of patients (2).

Despite this growing clinical deployment (4,5,8), most HTS efforts lack standardized infrastructure for data storage and analysis. Screening data are typically managed in *ad hoc* formats such as Excel spreadsheets, with analysis performed through one-off scripts or manual workflows that vary across institutions and even across projects within the same laboratory. A recent review of functional precision oncology highlighted that the heterogeneity of analytical approaches across programs remains a major barrier to cross-study comparison and clinical adoption (4). This fragmentation limits reproducibility, makes cross-cohort comparison difficult, and creates computational bottlenecks that ultimately slow the translation of drug screening results into clinical action. The gap is particularly acute in translational settings where screening results must be rapidly processed and communicated to clinicians, such as molecular tumor board meetings where drug sensitivity data inform time-sensitive treatment decisions for patients with limited therapeutic options.

Existing computational tools address individual aspects of the HTS workflow but have various limitations. Single-purpose metric calculators such as GRcalculator (9) and the DSS framework (10,11) compute specific dose-response metrics but do not handle data storage, multi-screen management, or downstream prioritization. More comprehensive platforms each address a different workflow aspect but leave critical gaps: ActivityBase (IDBS) (12) provides industrial-grade data management but is proprietary and designed for drug discovery rather than patient-centric workflows; Thunor (13) emphasizes interactive visualization but lacks cohort-level prioritization; HTSplotter (14) processes single screens end-to-end but without providing data storage solutions or cross-sample normalization; and iTReX (15) offers mono- and combination therapy analysis but operates per-screen without multi-sample data management. There is a need for an integrated, extendable, and open-source framework that combines standardized data storage, drug nomenclature standardization, cohort-level prioritization, custom metric extensibility, and automated reporting to support reproducible and scalable HTS analysis in precision oncology.

Here we present DS5, an open-source Python framework and an HDF5-based file format for standardized storage, quality control, dose-response analysis, drug prioritization, and automated reporting of HTS data. The framework is designed for any setting where drug screening data need to be stored, processed, and analyzed, from large-scale pharmacogenomics research to clinical precision oncology programs. Three design principles guide the architecture: ***reproducibility*** through raw data immutability, standardized drug nomenclature via RxNorm, and configurable quality control parameters with sensible defaults; ***scalability*** through HDF5’s disk-based architecture that handles datasets ranging from small clinical cohorts to hundreds of thousands of screens without requiring high-performance computing; and ***flexibility*** through an extensible design that supports user-defined custom metrics and analyses built on top of the core data structure. We validate DS5 against two large public pharmacogenomics datasets (GDSC2 (16,17) and CTRP (18,19)), demonstrate its flexibility with custom growth-rate adjusted metrics (20) on a published PDxO breast cancer dataset (21) with *in vivo* validation, and show real-world clinical application in an ongoing pediatric brain tumor precision oncology initiative where DS5 supports treatment discussions at molecular tumor board meetings.

## Results

### A unified framework for HTS data storage, analysis, and drug prioritization

Existing HTS tools each address one or two pieces of the screening workflow but force researchers to assemble heterogeneous pipelines from spreadsheets, single-purpose scripts, and proprietary platforms. DS5 is designed to fill this gap with a single open-source framework that integrates standardized storage, drug nomenclature standardization, reproducible quality control and dose-response analysis, cohort-level drug prioritization, and clinician-accessible reporting. **Figure 1A** illustrates the overall framework architecture; **Figure 1B** summarizes the end-to-end analytical workflow from raw plate readouts to patient-specific reports.

**Figure 1.**
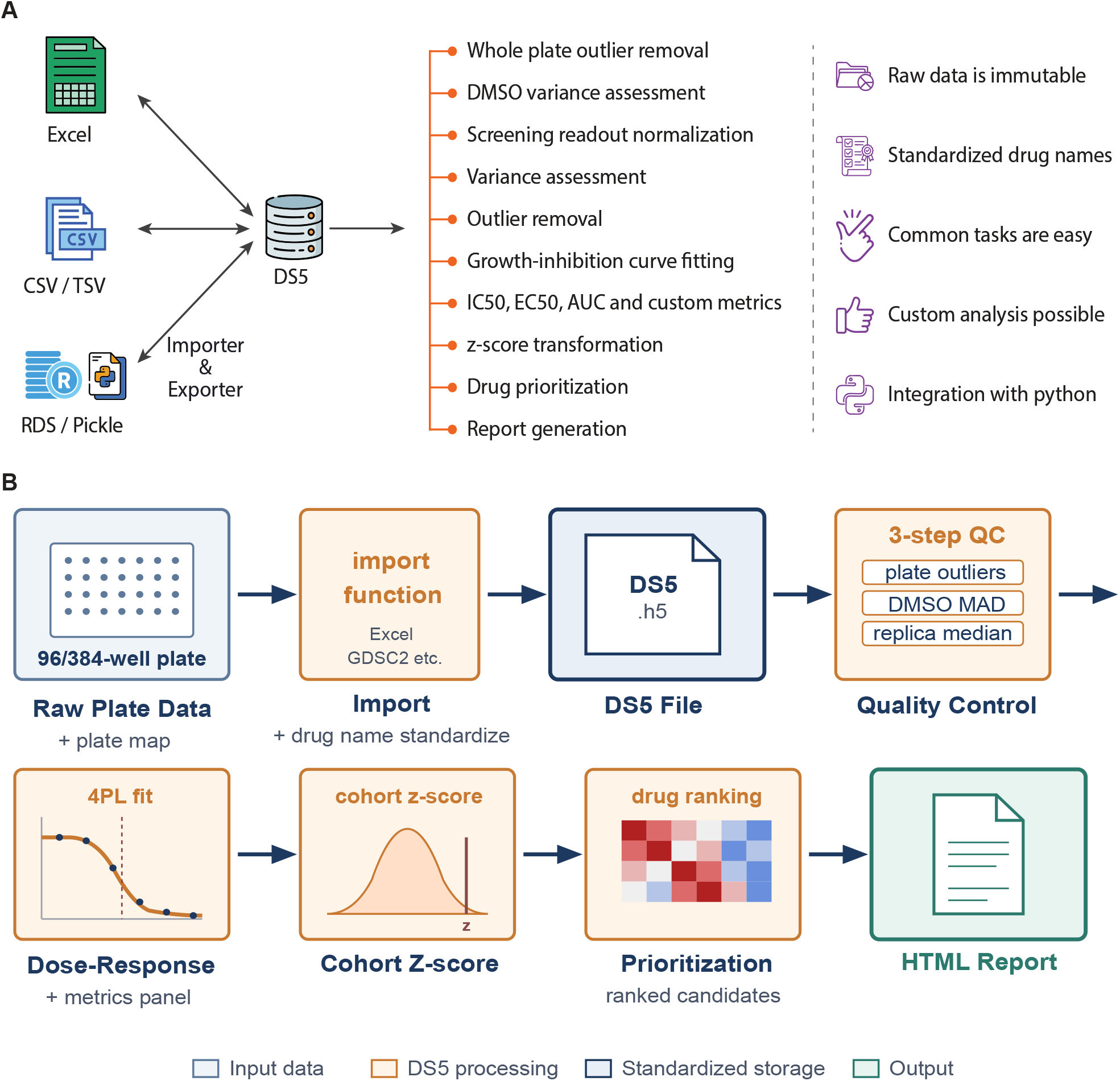
Overview of the DS5 Framework. **(A)** Schematic of the DS5 framework architecture, from data import through quality control, dose-response modeling, cohort-level drug prioritization, and automated reporting. **(B)** End-to-end analytical workflow from raw plate readouts to patient-specific reports.

### Standardized, immutable storage

Reproducibility in HTS analysis depends on preserving raw measurements. DS5 stores screening data in an HDF5-based hierarchical schema in which raw plate readouts are immutable, and every QC decision and analytical result is recorded as a separate attribute. This design provides full provenance from final metrics back to the original well-level measurements, supporting both scientific reproducibility and clinical auditing.

### Drug nomenclature standardization

Drug names vary widely across institutions, vendors, and screening libraries, creating a persistent barrier to cross-cohort comparison and to integration with clinical databases. On data import, DS5 resolves drug names against RxNorm and falls back to PubChem for compounds outside the clinical scope of RxNorm, assigning interoperable identifiers suitable for integration with clinical databases.

### Quality control

Plate-based HTS measurements are sensitive to dispensing errors, plate-edge effects, machine artifacts, and biological variation in baseline viability, all of which can distort downstream dose-response estimates. DS5 implements a configurable three-step QC pipeline consisting of plate-wide outlier removal, DMSO vehicle control filtering, and per-drug median-ratio replicate filtering, with all decisions logged within the DS5 file for retrospective inspection.

### Dose-response modeling and extensible metrics

DS5 fits a four-parameter logistic (4PL) model to normalized inhibition data and computes a comprehensive panel of pharmacological metrics (EC50, IC50, Emax, AUC, and three Drug Sensitivity Score variants-DSS1, DSS2 and DSS3; Table 1). DSS1 scores absolute drug sensitivity by integrating the area above a minimum-response threshold, DSS2 further normalizes by the tested concentration range to enable cross-drug comparison, and DSS3 additionally adjusts for the maximum achievable effect. Because the most informative metric depends on the experimental design and biological question, DS5 also exposes a custom-metric interface: any function operating on the standardized data structure can be registered and computed alongside the built-in measures, as we demonstrate later with growth-rate (GR) metrics on PDxO data.

**Table 1.**
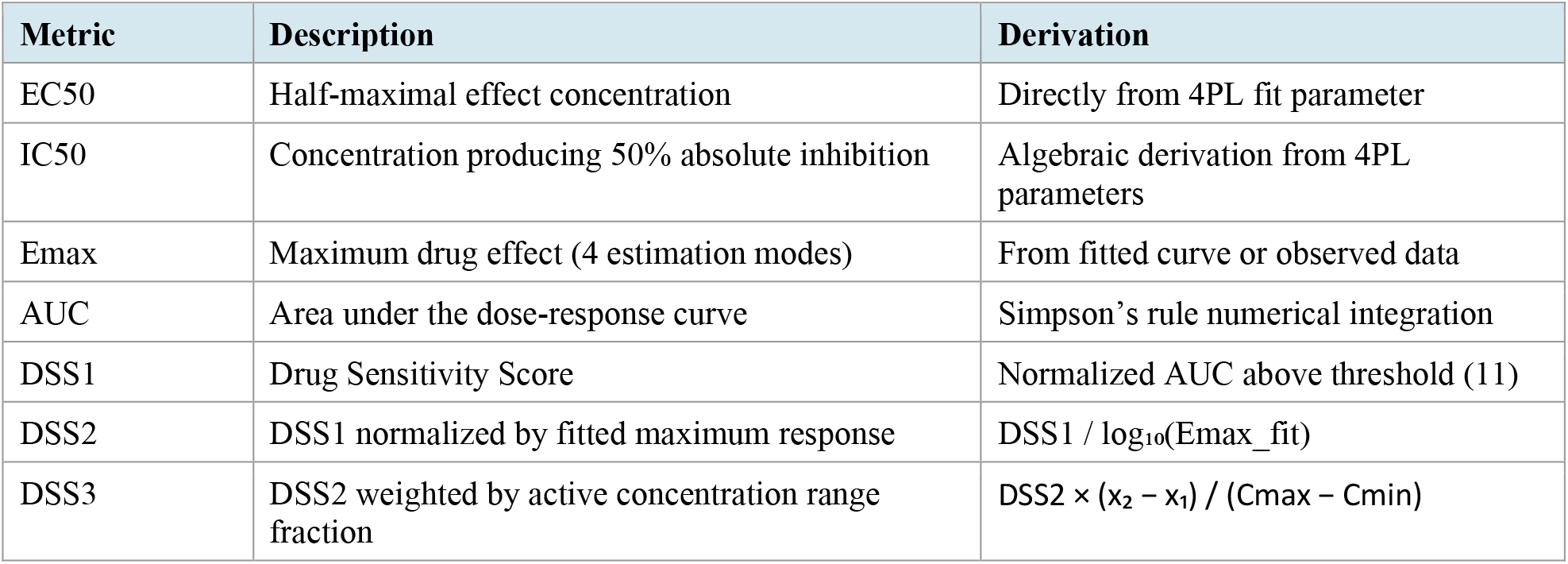
Drug Sensitivity Metrics Computed by DS5.

| Metric | Description | Derivation |
| --- | --- | --- |
| EC50 | Half-maximal effect concentration | Directly from 4PL fit parameter |
| IC50 | Concentration producing 50% absolute inhibition | Algebraic derivation from 4PL parameters |
| Emax | Maximum drug effect (4 estimation modes) | From fitted curve or observed data |
| AUC | Area under the dose-response curve | Simpson's rule numerical integration |
| DSS1 | Drug Sensitivity Score | Normalized AUC above threshold (11) |
| DSS2 | DSS1 normalized by fitted maximum response | $DSS1 / \log_{10}(Emax\_fit)$ |
| DSS3 | DSS2 weighted by active concentration range fraction | $DSS2 \times (x_2 - x_1) / (Cmax - Cmin)$ |

### Cohort-level drug prioritization

For clinical and translational applications, identifying drugs to which a patient’s tumor cells are exceptionally sensitive *relative to other patients* requires cohort-level normalization rather than absolute metric values alone. DS5 computes z-score transformations of any drug sensitivity metric (e.g., DSS, IC50, AUC, custom metrics) relative to a user-defined reference cohort and ranks drugs by their cohort-normalized scores. Cohort membership is defined at the screen level, allowing users to flexibly include or exclude individual screens. This design also accommodates situations in which a single patient contributes multiple screens, such as longitudinal samples or screens generated under different culture conditions, enabling construction of a reference distribution tailored to the specific analytical question.

### Automated reporting

Translating screening data into clinical action requires sharable, interpretable summaries. DS5 generates self-contained HTML reports that consolidate patient-specific prioritization with QC plots, cohort-level metric distributions, and dose-response curves into a single document suitable for molecular tumor board discussion or collaborator review.

### Quality Control Pipeline Demonstration

We illustrate the DS5 QC pipeline on clinical screening data from in-house patient-derived pediatric brain cancer cell lines. Because plate-based HTS measurements are prone to dispensing errors, well-position effects, and machine artifacts, removing aberrant data before normalization and curve fitting is essential to prevent these artifacts from propagating into downstream dose-response estimates. All three QC steps are demonstrated below.

For each screen experiment, percentile-based filtering identified and removed high and low intensity outlier wells likely to be machine measurement artifacts, reducing valid readouts from 384 to 344 in a representative screen (***Figure 2A***). MAD-based filtering of DMSO vehicle control wells confirmed tight baseline distributions across screens and flagged one outlier well in sample UW422 and sample UW422-2% (***Figure 2B***). For each individual drug-sample combination, the median-ratio filter flagged replicates deviating substantially from their dose-group peers. For example, 2 of 15 data points were flagged for Panobinostat in a single screen (***Figure 2C***). Removing these outliers before normalization and curve fitting prevents aberrant values from propagating into downstream dose-response estimates and drug sensitivity assessments. All QC decisions are logged within the DS5 file, providing the full provenance trail required for clinical auditing.

**Figure 2.**
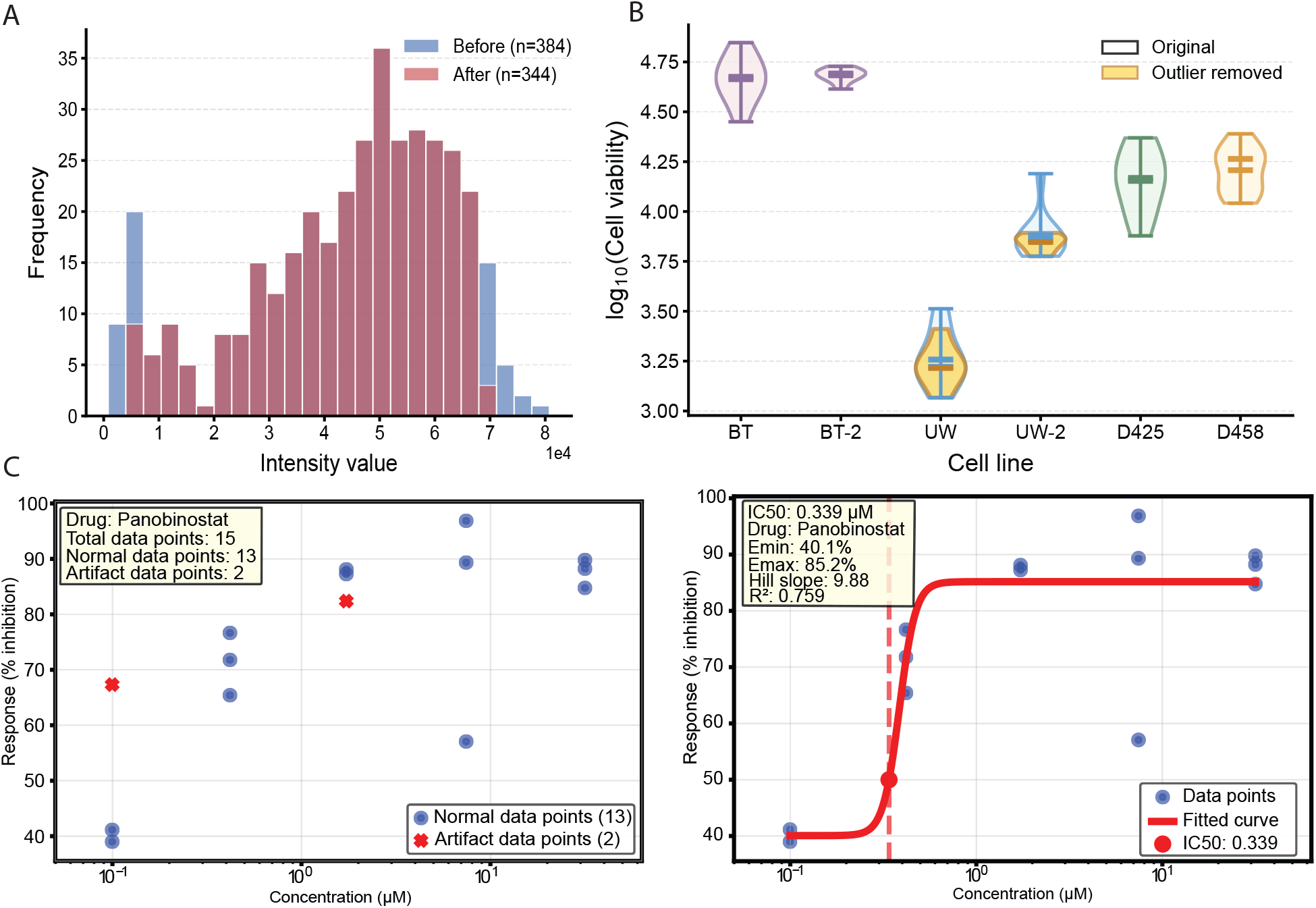
Quality Control Pipeline illustrated using patient-derived pediatric brain cancer cell lines data. **(A)** Screen-level quality control via percentile-based outlier removal. Histogram of raw CellTiter-Glo intensity values from a single drug screening plate (BT_test1) before (blue, n = 384) and after (red, n = 344) percentile-based filtering. High-intensity outlier wells, likely reflecting machine measurement artifacts, are removed to prevent downstream normalization artifacts. Data shown are from patient-derived pediatric brain cancer cell lines. **(B)** DMSO vehicle control quality control. Log_10_-transformed CellTiter-Glo signals from DMSO wells (n=9 per condition) across four patient-derived pediatric brain cancer cell lines cultured in stem cell media (BT12, UW422, D425, D458) or Advanced DMEM/F12 + 2% serum (BT12-2%, UW422-2%). Overlaid violins show data before and after outlier removal (z-score |z| > 3.5). One outlier removed from UW422 and UW422-2% each. **(C)** Drug-sample-level quality control via per-replicate median-ratio outlier detection. Left: scatter plot of % inhibition versus drug concentration (µM) for Panobinostat in a single screen (Sample UW422), with normal data points (blue, n = 13) and artifact replicates (red, n = 2) identified based on the median-ratio filter (ratio > 5 relative to group median at the same dose). Right: 4-parameter logistic (4PL) dose-response curve fitted to the filtered data.

### Validation with GDSC2 and CTRP Public Pharmacogenomics Data

To validate DS5’s analytical pipeline, we imported and analyzed two large independent pharmacogenomics datasets: the Genomics of Drug Sensitivity in Cancer (GDSC2) (16,17) which profiled ∼970 human cancer cell lines against ∼280 anticancer compounds, and the Cancer Therapeutics Response Portal version 2 (CTRPv2) (18,19,22) which profiled ∼860 cancer cell lines against ∼480 small-molecule probes and drugs. These datasets, processed through two independent published analytical pipelines, provide a rigorous benchmark for cross-validating DS5’s independently computed values.

GDSC2 raw dose-response data were imported directly into the DS5 format using the framework’s built-in import function. Although the GDSC consortium had performed initial curation and quality control prior to public release, we additionally applied DS5’s standard QC pipeline to ensure uniform processing across all downstream analyses. All metrics were recomputed through DS5’s standard pipeline. To illustrate DS5’s built-in visualizations at two levels of granularity, ***Figure 3A*** shows a within-screen inhibition heatmap of DS5-computed values across selected drugs and concentrations for a single cell line (Daoy_7230). ***Figure 3B*** shows a DSS1 heatmap across a user-selected subset of drugs and cell lines for cohort-level cross-sample comparison. Together, these visualizations enable researchers to identify dose response patterns at the single experiment level and to compare drug sensitivity across samples efficiently.

**Figure 3.**
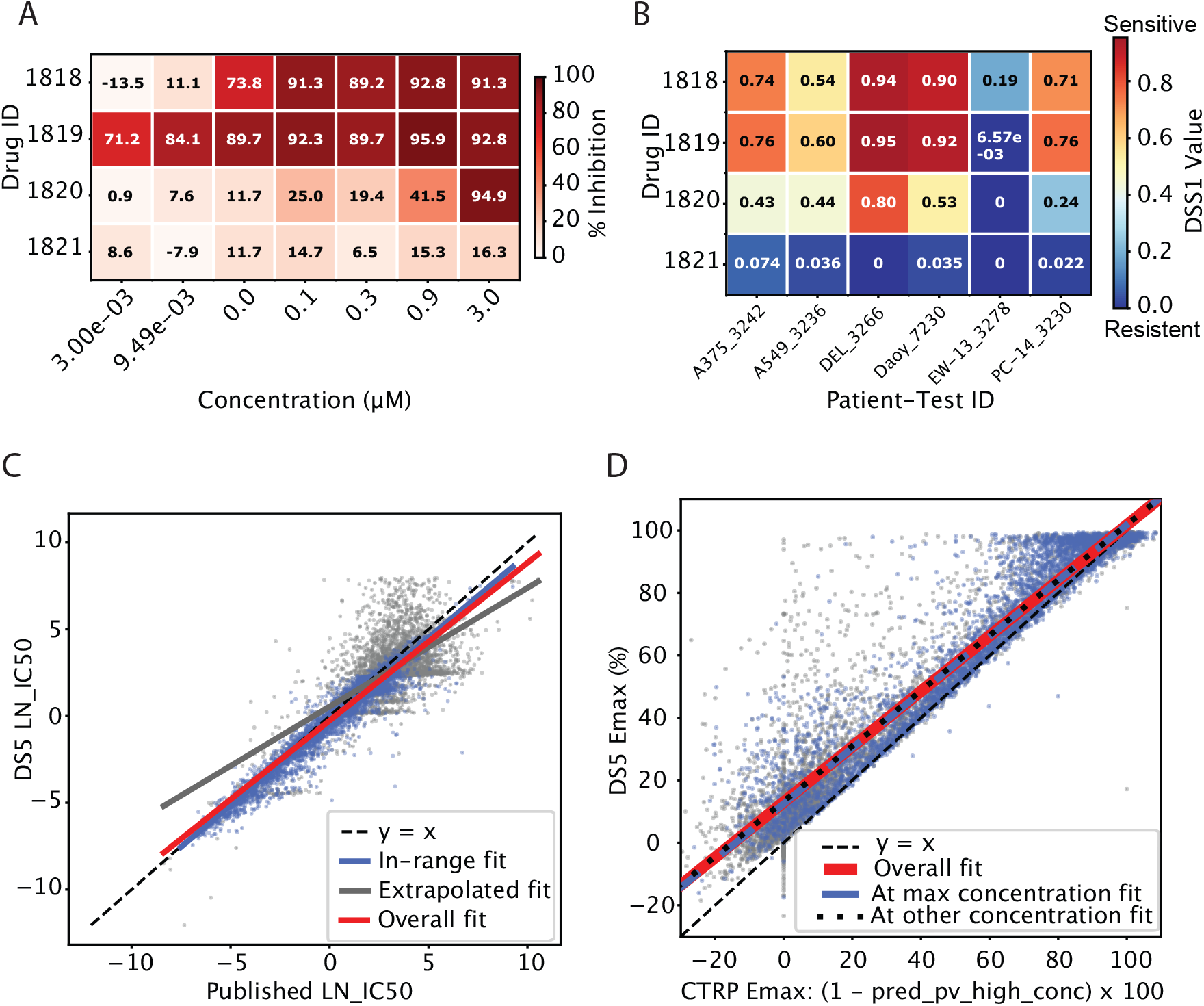
Validation of DS5 against published GDSC2 and CTRPv2 pharmacogenomics datasets. **(A)** Within-screen inhibition heatmap of DS5-computed percent inhibition values for four selected drugs (drug IDs 1818–1821) across seven concentrations (3.0e-03 to 1.0 µM) in the GDSC2 Daoy_7230 screen, illustrating DS5’s built-in visualization for per-screen dose-response exploration. **(B)** DSS1 heatmap for a user-selected subset of four drugs across six GDSC2 cell line screens (A375, A549, DEL, Daoy, EW-13, PC-14), demonstrating the framework’s flexibility in subsetting cohort data for focused cross-screen comparison. **(C)** Concordance between DS5-computed and GDSC2-published natural-log IC50 (LN_IC50) values across 102,637 drug-cell line pairs (10,000 randomly sampled pairs shown). Blue points indicate pairs where the fitted IC50 fell within the experimentally tested concentration range (Pearson r = 0.973, n = 65,749); gray points indicate pairs requiring extrapolation beyond the tested doses (Pearson r = 0.654, n = 36,888). Overall Pearson r = 0.918. Dashed line: y = x. **(D)** Concordance between DS5-computed and CTRPv2-published Emax values across 326,305 measurements (10,000 randomly sampled points shown). Blue points indicate cell-line / drug pairs for which maximum inhibition is achieved at the maximum tested concentration (Pearson r = 0.961, n = 174,661); gray points indicate cell-line /drug pairs for which maximum inhibition is achieved at non-maximum concentrations (Pearson r = 0.938, n = 151,644). Overall Pearson r = 0.955. Dashed line: y = x.

Quantitative validation against published natural-log IC50 values (LN_IC50) showed strong agreement across 102,637 drug-cell line pairs (Pearson r = 0.918; ***Figure 3C***). To characterize this concordance more precisely, we distinguished pairs where the fitted IC50 fell within the tested concentration range (n = 65,749) from those requiring extrapolation beyond the tested doses (n = 36,888). Both subsets showed high correlation with published values, with the in-range subset achieving slightly higher concordance (Pearson r=0.973) — consistent with the expectation that fitted estimates are more reliable when the IC50 falls within the tested concentration window.

For CTRPv2, we used Emax (maximum drug effect) as a complementary validation metric that directly tests the fit at the maximum effect, orthogonal to the IC50 validation in GDSC2. DS5- computed Emax values were highly concordant with CTRP-published values across 326,305 measurements (Pearson r = 0.955; ***Figure 3D***), with tighter agreement for measurements at the maximum tested concentration (Pearson r=0.961, n = 174,661) than at other concentrations (Pearson r=0.938, n = 151,644).

Together, the strong quantitative agreement across IC50 (GDSC2; n = 102,637) and Emax (CTRPv2; n = 326,305), drawn from two independent pharmacogenomics pipelines, establishes the analytical reproducibility of DS5’s core metric calculations.

### Custom GR Metric Implementation and In Vivo–Validated Drug Prioritization on Breast Cancer PDxO Data

To demonstrate DS5’s flexibility beyond its built-in metrics, we imported a published patient-derived xenograft organoid (PDxO) breast cancer drug screening dataset (21) comprising 45 compounds tested across 16 PDxO models at 8-point dose ranges with 3 biological × 4 technical replicates per condition (***Figure 4A***). The original study employed growth rate (GR) metrics, which account for differences in cell division rates and are not natively built into DS5’s standard pipeline. We implemented GR-based analysis as a custom extension on top of DS5’s data structure, illustrating how users can adapt the platform to specialized analytical needs without modifying the core package.

**Figure 4.**
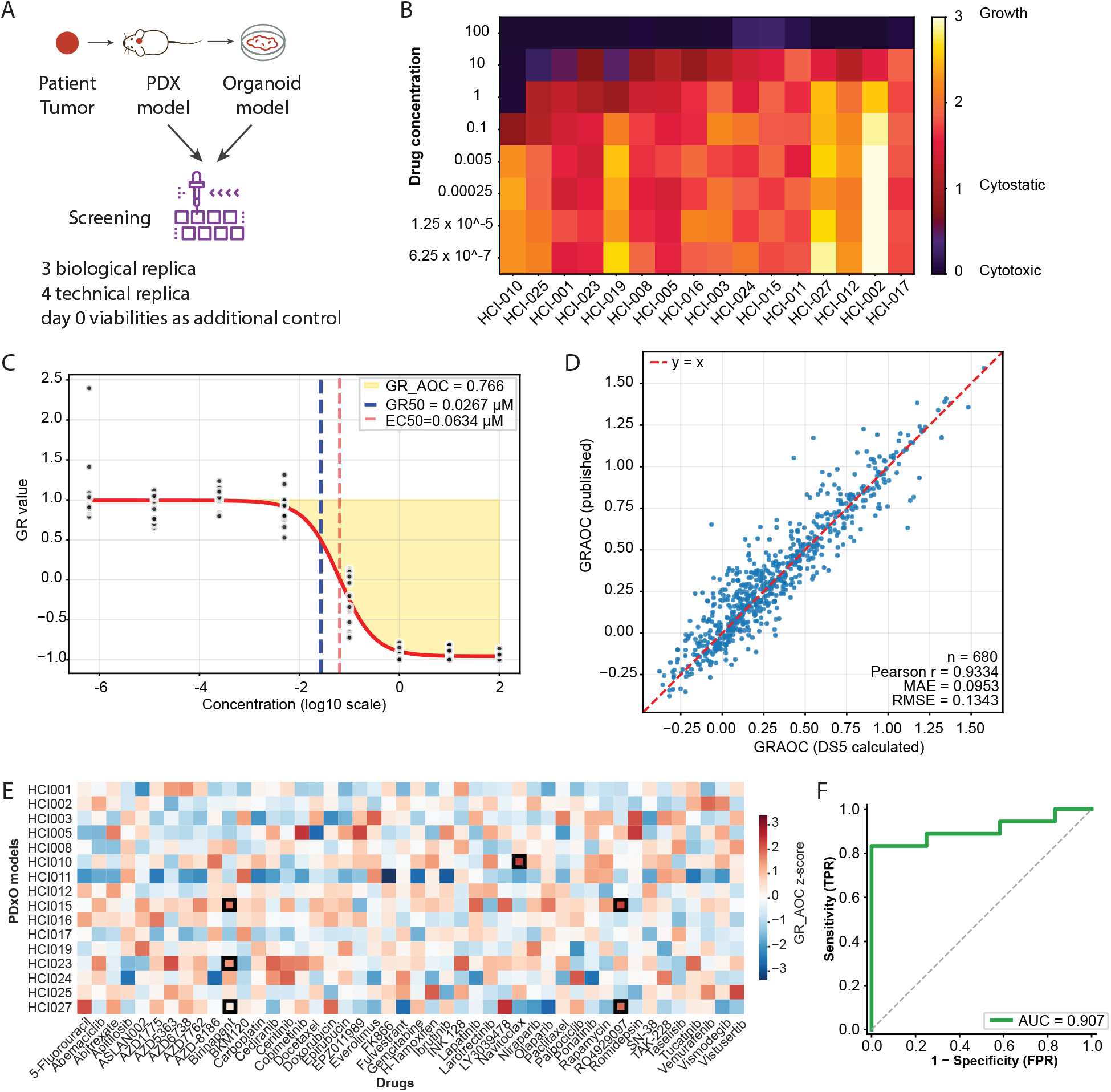
Reproduction and Extension of PDxO Breast Cancer Drug Screening Analysis. **(A)** Schematic of the PDxO drug screening workflow. Patient tumor-derived PDX models were used to establish organoid cultures (PDxOs), which were screened against a panel of 45 compounds in an 8-point dose-response format with 3 biological replicates and 4 technical replicates per condition. Day-0 viability measurements were included as an additional normalization control to account for variable organoid growth rates across models. **(B)** DS5 reproduction of drug response profiles for navitoclax. Heatmap displays GR across an 8-point dose range (y-axis) and PDxO models (x-axis). Results are concordant with published drug response profiles reported in Guillen et al., 2022. **(C)** GR dose-response curve for navitoclax in HCI-010 fitted by DS5. The y-axis displays growth rate-adjusted estimates from CellTiter-Glo 3D (CTG-3D) cell viability assays. The x-axis shows log10-transformed drug concentrations across an 8-point dose range. Each dot represents one of 12 replicates (3 biological replicates and 4 technical replicates each). Annotations include EC50, GR50, and GR_AOC. **(D)** Scatter plot of DS5-calculated versus published GR_AOC values across all drug-model pairs (n = 680). Strong agreement between DS5- derived and reported values demonstrates that DS5 accurately reproduces GR_AOC estimates across the full dataset. **(E)** GR_AOC z-score heatmap of drug response profiles across 16 models. Z-scores for each drug were calculated relative to the full PDxO cohort, with red indicating relative sensitivity and blue indicating relative resistance. Black borders highlight drug-model pairs with confirmed *in vivo* sensitivity reported in Guillen et al., 2022 (Navitoclax in HCI-010; Birinapant in HCI-015, HCI-023, and HCI-027; and RO4929097 in HCI-015 and HCI-027), demonstrating its utility for drug prioritization in precision oncology settings. **(F)** ROC analysis of *in vitro* drug sensitivity as a predictor of *in vivo* response. GR AOC z-scores from PDxO drug screening were used to predict *in vivo* tumor response across 30 drug–PDX model pairs (5 drugs, multiple HCI models). *In vivo* responders were defined as models with >40% tumor growth reduction. The ROC curve was constructed by sweeping the GR AOCz threshold and computing sensitivity and specificity at each value. An AUC of 0.907 indicates that *in vitro* GR AOCz is a strong predictor of *in vivo* drug response.

Using the custom GR implementation, DS5 reproduced the original findings of Guillen et al. with high fidelity. A heatmap of GR values for navitoclax across the dose range and PDxO models (***Figure 4B***) visually matched the published profiles. At the individual drug level, DS5’s dose-response fitting accurately captured the GR curve for navitoclax in HCI-010 (***Figure 4C***). A direct comparison of DS5-calculated versus published GR_AOC values across all 680 drug-model pairs showed strong quantitative agreement (***Figure 4D***).

Beyond reproducing published metrics, we applied DS5’s cohort-level drug prioritization to identify PDxO models with exceptional drug sensitivity. For each drug, GR_AOC values were z-transformed across all 16 models, so that a higher z-score indicates greater sensitivity relative to the cohort. The resulting GR_AOC z-score heatmap (**Figure 4E**) revealed that drug-model pairs with confirmed in vivo sensitivity reported in Guillen et al., including navitoclax in HCI-010, birinapant in HCI-015/023/027, and RO4929097 in HCI-015 and HCI-027, consistently ranked among the highest z-scores within their respective drug columns. ROC analysis using in vitro GR_AOC z-scores to predict in vivo tumor response across 30 drug-PDX pairs yielded an AUC of 0.907 (**Figure 4F**), demonstrating that DS5’s cohort-level prioritization can recover drug sensitivities with translational relevance.

### Clinical Application: Drug Prioritization for Preclinical Validation in Pediatric Brain Cancer Initiative

To demonstrate DS5’s utility in a real-world preclinical drug development setting, we applied the framework to an ongoing in-house pediatric brain cancer precision oncology initiative with a functional screening component. At the current stage of the pilot study, cell-line based *in vitro* drug sensitivity screening is used to prioritize candidate drugs for downstream PDX-based *in vivo* validation. We screened four patient-derived pediatric brain cancer cell lines (BT12, UW422, D425, D458) against a panel of 25 compounds at 5 concentrations. Two cell lines (BT12 and UW422) were additionally screened in Advanced DMEM/F12 + 2% serum alongside the standard stem cell media condition to assess condition-dependent drug response, yielding 6 screens in total. DS5’s three-step QC pipeline was applied to all six screens prior to analysis (representative QC examples shown in ***Figure 2***).

Plate-level visualization of drug responses for BT12 across the two culture conditions (***Figure 5A***) revealed largely consistent sensitivity profiles between stem cell media and 2% serum, supporting the biological robustness of the observed responses.

**Figure 5.**
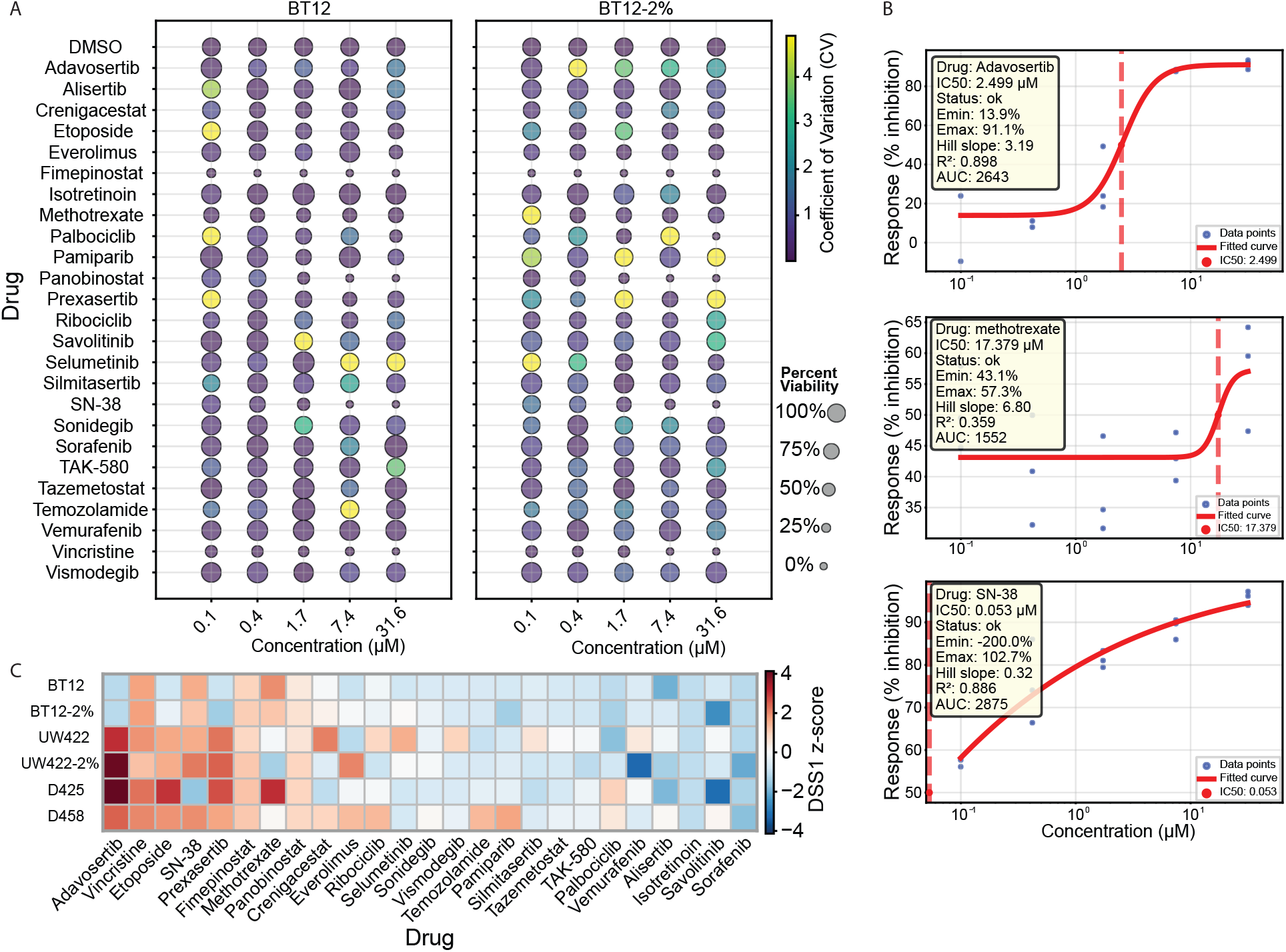
Clinical Application in Pediatric Brain Tumor Precision Oncology. **(A)** Plate-level drug response patterns. BT12 cells cultured in stem cell media (left) and 2% serum (right) show consistent drug responses across 25 compounds and five concentrations. Bubble size = viability; color = CV across replicates. Drug response patterns are largely similar between conditions. **(B)** Dose-response curves for top-ranked drug–cell line pairs. Curves shown for three drug–cell line combinations identified as exceptional responders in panel (C): Adavosertib in UW422-2% (left), Methotrexate in D425 (center), and SN-38 in D458 (right). Each curve displays replicate data points with four-parameter logistic fit; IC50 values are indicated by dashed red lines. The three pairs illustrate distinct response profiles: Adavosertib shows a classic sigmoidal response with moderate potency, Methotrexate exhibits a steep transition at higher concentrations, and SN-38 demonstrates high potency at sub-micromolar concentrations. These candidates were selected for further clinical consideration based on their cohort-normalized sensitivity rankings.**(C)** Drug prioritization via patient cohort normalization. DSS1 z-scores relative to 15 patient-derived pediatric brain cancer cell lines. Z-scores were calculated using cohort mean and standard deviation for each drug. (left = high sensitivity, right = resistance). Exceptional responses identified for Adavosertib in UW422/D425, Vincristine across cell lines, and Methotrexate for D425. Cell line-specific patterns reveal distinct therapeutic vulnerabilities.

To prioritize drug candidates for in vivo testing, we computed DSS1 z-scores for each cell line relative to a reference cohort of 15 patient-derived pediatric brain cancer samples from an ongoing parallel program, used here as a normalization background (***Figure 5C***). This cohort-normalized ranking transforms absolute sensitivity values into relative scores, identifying drugs that produce exceptional sensitivity in specific cell lines compared to the broader patient population. Such cell-line-selective patterns are most informative for selecting candidate combinations for resource-intensive *in vivo* experiments. Notably, the same compound can show significantly different selectivity across different cell lines. For example, Methotrexate ranked among the top-sensitive drugs in BT12 and D425, but fell outside the top 4 for UW422 and D458, illustrating how cohort normalization reveals cell-line-specific therapeutic vulnerabilities that absolute sensitivity scores alone would miss. Based on this prioritization, both top-sensitive and top-resistant drugs were selected for each cell line (***Table 2***): top-sensitive drugs as candidate therapeutic hits, and top-resistant drugs as built-in negative controls to validate that the *in vivo* assay can distinguish responders from non-responders. These selections are currently advancing toward *in vivo* validation experiments within the pilot study.

**Table 2.** Drug candidates selected for in vivo validation based on DS5 cohort-level prioritization.

| Cell line | Top-sensitive drugs | Top-resistant drugs |
| --- | --- | --- |
| BT12 | Methotrexate, SN-38, Vincristine, Fimepinostat | Pamiparib, Sorafenib, Tovorafenib |
| UW422 | Prexasertib, Adavosertib, SN-38, Vincristine | Everolimus, Ribociclib, Sorafenib |
| D425 | Methotrexate, Adavosertib, Etoposide, Prexasertib | Selumetinib, Tovorafenib, SN-38 |
| D458 | Ribociclib, Pamiparib, Adavosertib, SN-38 | Isotretinoin, Selumetinib, Silmitasertib |

For the top-ranked drug-cell line pairs, DS5 generated dose-response curves for visual inspection of drug responses (***Figure 5B***). The three highlighted curves (Adavosertib in UW422-2%, Methotrexate in D425, and SN-38 in D458) show distinct response profiles, with classical sigmoidal kinetics, steep high-dose transitions, and sub-micromolar potency, respectively.

## Discussion

DS5 addresses a critical gap in the HTS tool landscape: a unified, lightweight framework that combines standardized data storage, reproducible analysis, and clinician-accessible reporting in a single open-source package. Validation against GDSC2 and CTRPv2 establishes analytical accuracy at scale, with strong concordance for both IC50 (Pearson r = 0.973) and Emax (r = 0.961) across more than 428,000 drug-cell line measurements. The PDxO breast cancer analysis demonstrates extensibility through user-defined GR metrics and shows that DS5-based drug prioritization recovers in vivo–validated drug sensitivities (ROC AUC = 0.907). The pediatric brain cancer initiative demonstrates real-world translational utility, with DS5-driven prioritization currently informing *in vivo* validation experiments in an active pilot program.

As functional drug screening expands from research laboratories into clinical decision-making (1–4,23,24), the requirements for analytical infrastructure shift substantially. Clinical and translational applications demand not only accurate analytics but also data provenance, audit trails, standardized nomenclature for integration with clinical databases, and reporting formats accessible to non-computational users. DS5’s design directly addresses these translational requirements through raw data immutability for provenance, RxNorm/PubChem standardization for cross-institutional interoperability, and automated HTML reporting for clinician accessibility. The framework’s design also aligns with FAIR principles (25) through standardized HDF5 schemas with embedded metadata (Findability, Accessibility), drug nomenclature standardization (Interoperability), and raw data preservation alongside documented analytical provenance (Reusability).

A concrete example of the value of standardized infrastructure arose during clinical use. An initial observation that two screens with markedly different DMSO control signals appeared to differ in drug response variability prompted a cohort-wide investigation of whether baseline cell viability systematically affects sensitivity measurements. Using DS5’s cohort-level query capabilities, we computed per-drug inhibition variability across all screens ordered by DMSO viability and found no systematic relationship **(Supplementary Figure 1**), suggesting that normalization adequately corrects for baseline variation across heterogeneous patient-derived samples. Such *ad hoc* cohort-wide hypothesis testing is only practical when data from multiple patients are stored in a consistent, queryable format. This is precisely the infrastructure that DS5 provides and that is absent from workflows based on disconnected spreadsheets.

Several limitations of the current framework should be noted, each pointing toward planned development priorities. First, DS5 currently requires plate-map-based Excel input; supporting long (tidy) format with configurable column mappings would further reduce preprocessing burden for users importing from diverse screening platforms. Second, DS5 operates through

Python scripting and Jupyter notebooks, limiting accessibility for users without coding experience. A planned web-based interface would enable bench scientists to upload data, run analyses, and explore results interactively without writing code. Third, the framework currently supports only single-agent dose-response analysis and does not handle combination screens, which require synergy scoring and dose-response surface modeling. Extending DS5 to support combination analysis would address the growing importance of combination therapies in precision oncology (26). The open-source, modular architecture of DS5 is designed to facilitate community contributions addressing these and other extensions.

In summary, DS5 provides a comprehensive, open-source framework for HTS data management and analysis that fills a critical unmet need in the field. Its demonstrated accuracy across large public pharmacogenomics datasets, flexibility in accommodating custom metrics, and ongoing deployment in an active preclinical drug development program together demonstrate that DS5 is ready for broader adoption. As functional drug screening continues to expand into clinical practice, standardized computational infrastructure like DS5 will be essential for ensuring that screening data are analyzed reproducibly, interpreted consistently, and communicated effectively to support patient care.

### Data and Code Availability

DS5 is freely available on the Python Package Index (PyPI) and can be installed via “pip install DS5”. Source code, comprehensive documentation, example Jupyter notebooks, and mock datasets are available on GitLab at [https://gitlab.com/qiao-lab/ds5]. The GDSC2 and CTRP datasets analyzed in this study are publicly available from their respective repositories (16–19,22). The PDxO breast cancer dataset was obtained from Guillen et al. (21). Screening data from the cell-lines tested within the pediatric brain tumor initiative are available as **Supplemental Data**.

## Materials and Methods

### DS5 File Architecture and Data Storage

The DS5 format is built on HDF5 (27) and organizes screening data hierarchically as Patients → patient_id → test_id → (data table + plate map). Each screen is stored as two paired tables with matching dimensions: a raw readout table and a plate map encoding drug assignments, concentrations, and control positions. Raw data are stored read-only. All QC decisions and analytical results are written as separate attributes within the HDF5 hierarchy.

### RxNorm Drug Name Standardization

Drug names are resolved on import via a two-tier scheme: names are first queried against RxNorm (28), unmatched names are then queried against PubChem (29). Compounds not found in either database are flagged with a warning and stored under their raw names, so non-standard compounds do not block the import workflow.

### Data Import and Export

Users prepare raw screening data in the standard two-table format (data table and plate map with matching dimensions) and invoke an importer function to convert them to DS5. An Excel implementation is provided, as well as a dedicated importer for GDSC2 published dataset. The framework handles all downstream storage, organization, and analysis from that point. Processed metrics and normalized data can be exported back to Excel for downstream use.

### Quality Control

DS5 implements a configurable three-step sequential QC pipeline: (1) percentile-based plate-wide outlier removal, (2) MAD-based DMSO vehicle control filtering, and (3) per-drug median-ratio filtering to flag aberrant replicates within each dose group. All thresholds are user-configurable via a settings file. After outlier removal, drug well values are normalized to DMSO controls as percent inhibition. All QC decisions are logged within the DS5 file, and DS5 generates before/after QC comparison plots at the DMSO, per-sample drug response level, and plate level.

### Dose-Response Modeling and Sensitivity Metrics

Dose-response analysis in DS5 centers on a four-parameter logistic (4PL) model fitted to normalized percent inhibition data. From this fit, DS5 computes EC50, IC50, Emax, AUC, and three Drug Sensitivity Score variants (DSS1, DSS2, DSS3) (10,11). The DSS variants implement progressive normalization layers (DSS1: AUC above an activity threshold; DSS2: DSS1 divided by log_10_(Emax_fit); DSS3: DSS2 weighted by the active concentration-range fraction).

Definitions are summarized in Table 1. User-defined custom metrics can be registered and computed alongside built-in measures.

### Cohort-Level Analysis and Drug Prioritization

DS5 computes z-score transformations of any drug sensitivity metric (e.g., DSS, IC50, AUC, or custom metrics) relative to the distribution across a user-defined reference cohort. Cohort is defined per screen. Users may include or exclude any subset of screens (including multiple screens from the same individual). Drugs are then ranked by cohort-normalized score. Cross-patient and cross-drug comparisons are provided through integrated heatmap visualizations and ranked metric tables.

### Automated Report Generation

DS5 includes a Jupyter Notebook demonstrating the complete analytical workflow from data import through quality control, metric calculation, key visualization methods, and cohort-level drug prioritization. A dedicated reporting function additionally generates self-contained HTML reports that consolidate the resulting patient-specific drug prioritization with supporting evidence including QC plots, cohort-level metric distributions, and z-score-based prioritization plots into a single shareable document suitable for clinical discussions.

### Software Implementation and Installation

DS5 is distributed as an open-source Python package on the Python Package Index (PyPI) and can be installed with a single command (pip install DS5). The package runs on Python 3.11 or later and has minimal dependencies, all of which are automatically resolved during installation. The repository includes example Jupyter notebooks with a mock dataset that allows new users to execute the complete workflow without requiring their own screening data. The test suite, implemented with pytest, covers all core modules (data import, QC filtering, dose-response fitting, metric calculation, cohort-level prioritization, and report generation), achieving 86% line coverage. Source code, documentation, and example data are maintained on GitLab under the MIT license.

## Author Contributions

**Y.Q**. conceived, supported, and oversaw the development of the DS5 project. **J.L**. developed the initial prototype of many features in DS5. **H.Y**. further developed DS5 to completion, as well as performed data analysis, results interpretation, and manuscript preparation. **X.H**. and **G.M**. contributed to ideas, discussions, data generation, and data analysis. **C-H.Y**., **A.W**., and **S.C**. contributed to discussions and data generation. **P.M**. performed the drug screening experiments on the pediatric brain cancer patient derived cell-lines, and contributed to project discussions. All authors contributed to the preparation and editing of this manuscript.

## Supplementary Figures

**Supplementary Figure 1.**
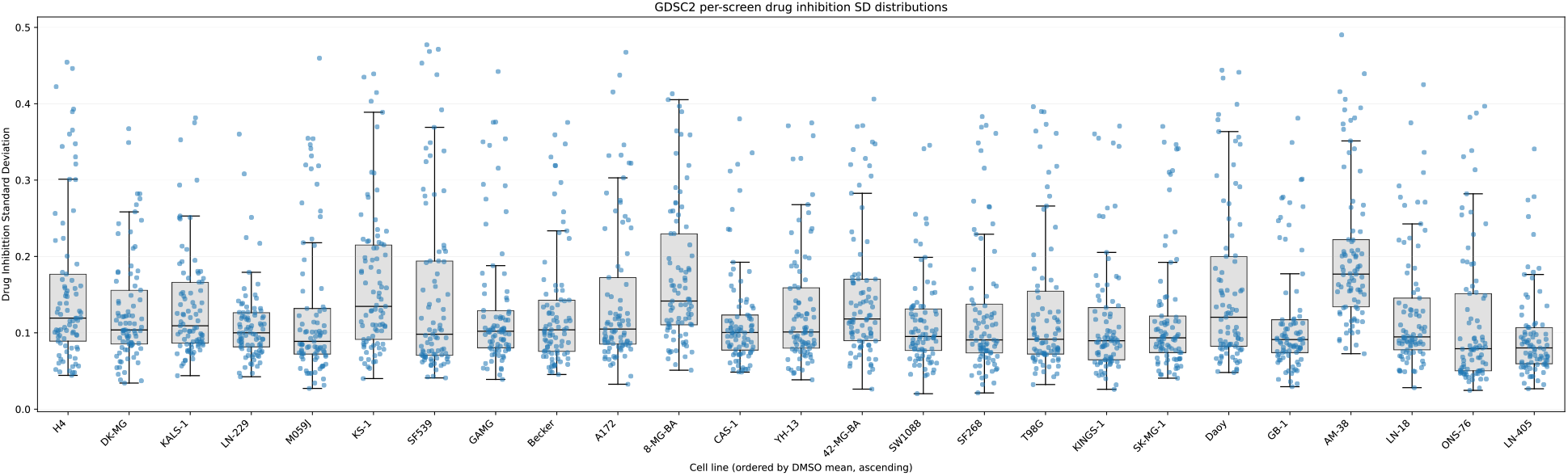
Drug response variability is independent of baseline DMSO viability across screens. For each screen, drug inhibition variability was quantified as the standard deviation of inhibition values across all tested compounds. Screens were ordered by mean DMSO control viability, and distributions of per-drug variability are shown as boxplots. Each point represents an individual drug measurement. No systematic association between baseline viability and drug response variability was observed across the cohort, indicating that normalization effectively compensates for baseline differences across heterogeneous samples.

